# The genetics of gene expression in a *C. elegans* multi parental recombinant inbred line population

**DOI:** 10.1101/2021.03.04.433879

**Authors:** Basten L. Snoek, Mark G. Sterken, Harm Nijveen, Rita J.M. Volkers, Joost Riksen, Philip C. Rosenstiel, Hinrich Schulenburg, Jan E. Kammenga

## Abstract

Studying genetic variation of gene expression provides a powerful way to unravel the molecular components underlying complex traits. Expression QTL studies have been performed in several different model species, yet most of these linkage studies have been based on genetic segregation of two parental alleles. Recently we developed a multi-parental segregating population of 200 recombinant inbred lines (mpRILs) derived from four wild isolates (JU1511, JU1926, JU1931 and JU1941) in the nematode *Caenorhabditis elegans*. We used RNA-seq to investigate how multiple alleles affect gene expression in these mpRILs. We found 1,789 genes differentially expressed between the parental lines. Transgression, expression beyond any of the parental lines in the mpRILs, was found for 7,896 genes. For expression QTL mapping almost 9,000 SNPs were available. By combining these SNPs and the RNA-seq profiles of the mpRILs, we detected almost 6,800 eQTLs. Most *trans*-eQTLs (63%) co-locate in six newly identified *trans*-bands. The *trans*-eQTLs found in previous 2-parental allele eQTL experiments and this study showed some overlap (17.5%- 46.8%), highlighting on the one hand that a large group of genes is affected by polymorphic regulators across populations and conditions, on the other hand it shows that the mpRIL population allows identification of novel gene expression regulatory loci. Taken together, the analysis of our mpRIL population provides a more refined insight into *C. elegans* complex trait genetics and eQTLs in general, as well as a starting point to further test and develop advanced statistical models for detection of multi-allelic eQTLs and systems genetics studying the genotype-phenotype relationship.

## Introduction

Investigation of the genotype-phenotype relationship is at the heart of genetic research. The detection and description of allelic variants and genetic mechanisms have been a demanding task due to the quantitative nature of most phenotypic variation. Quantitative trait locus (QTL) mapping has been one of the methods of choice for finding the loci on which these allelic variants can be found. Many functional polymorphisms in plants and animals, including many model species such as model nematode *C. elegans*, have been discovered using QTL mapping [1-25]. Over the last decade molecular phenotypes such as transcript levels, protein levels and metabolites have also been used in QTL mapping [26-32]. Heritable variation in these molecular phenotypes often plays a role in heritable phenotypic variation [10, 30, 33]. Mapping expression QTLs (eQTLs) can provide insight into the transcriptional architecture of complex traits and have been conducted in model species such as *Arabidopsis thaliana* and *C. elegans* as well as several other taxa [26, 28-31, 34-41].

Most eQTL studies have been done on populations of recombinant inbred lines (RILs) originating from a cross between two different parental genotypes [26, 28-31, 34-40]. Inclusion of more than two parents can capture more genetic variation, increasing the number of detected QTLs, potentially allowing more precise mapping and therefore reducing the number of potential candidate causal genes to be verified [42]. Such a strategy was first used for *Arabidopsis* by developing a Multiparent Advanced Generation Inter-Cross (MAGIC) lines population consisting of 527 RILs developed from 19 different parental accessions [43]. Several other MAGIC populations have been developed since then for a range of species, including *C. elegans* [44-46].

Recently multi parental RIL (mpRILs) populations have been developed in *C. elegans* [45, 46]. These populations have been created using other strains than the most frequently used N2 strain and the Hawaiian CB4856 strain [26-31, 37, 47-61]. In this study we used the population of 200 mpRILs, derived from an advanced cross between four wild-types: JU1511 and JU1941 isolated from Orsay (France) and JU1926 and JU1931 isolated from Santeuil (France) (kindly provided by MA Félix, Paris, France; [45, 62]). In a previous study, the RNA-seq data of these mpRILs was used to obtain almost 9,000 SNPs variable between the four parental genotypes and used to identify QTLs for life-history traits [45]. The RNA was sampled from the mpRILs grown under standardised conditions (24°C, OP50, 48h after bleaching) and obtained from animals from two 6-cm dishes, with one RNA-seq replicate per mpRIL and two per parental isolate. To investigate the effect of multiple genetic backgrounds on gene expression, we used the RNA-seq data to associate gene expression levels to genetic variants present in the population. We compared the gene expression level differences between the parental wild isolates, calculated transgression, as well as heritability and mapped eQTLs. We identified six *trans*-bands, hotspots at which many *trans*-eQTLs co-locate, which we further studied by gene ontology enrichment. Lastly, we compared the eQTLs found in this study to the eQTLs found in previous eQTL studies in *C. elegans* [26, 28, 30, 31, 37, 39]. Together these results present the first insights into the genetic architecture of gene expression in a *C. elegans* multi parental RIL population.

## Results

### Gene expression differences between the parental lines

To study the effect of genetic variation on gene expression we used RNA-seq on a population of 200 multi parental recombinant inbred lines (mpRILs) [45], made from a cross between four parental lines isolated from Orsay, France (JU1511, JU1941) and Santeuil, France (JU1926, JU1931) [62]. The animals used were grown on two 6-cm dishes (24°C, OP50, 48h after bleaching) per sample pooled for RNA isolation, with one RNA-seq replicate per mpRIL and two per parental isolate. First, we determined the expression differences between the parental lines (**Supplement table 1**). Of the 12,029 detected transcripts we found 1,789 genes differently expressed between at least one parental pair (TukeyHSD p < 0.001; FDR < 0.05; **Figure 1**). Of the four strains, JU1926 was most different when compared to the other lines, with 409 genes being differently expressed between JU1926 and the other three lines. Thereafter, JU1941 was most different from the remaining two lines. These differences in gene expression between the parental lines are likely genotype dependent.

**Table 1:**
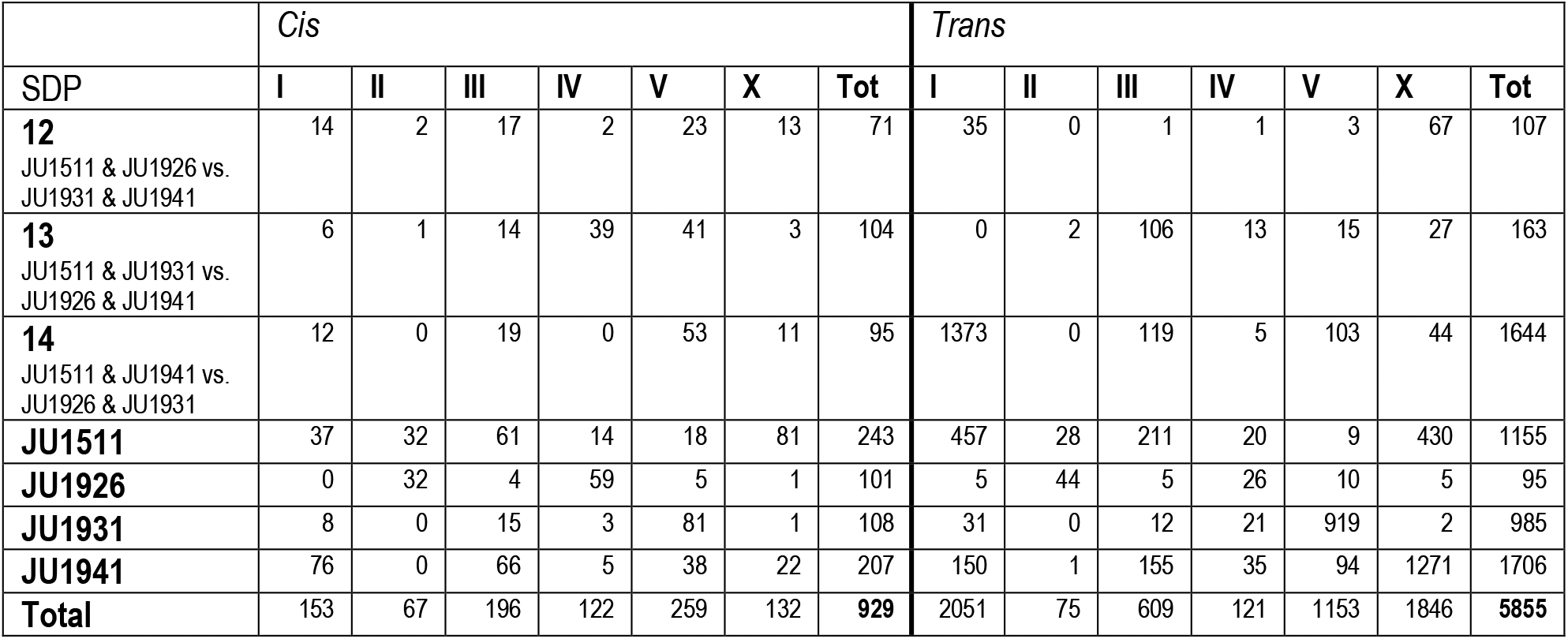
eQTLs per type (*cis*/*trans*) per chromosome per SNP Distribution Pattern (SDP).

|  | *Cis* |  |  |  |  |  |  | *Trans* |  |  |  |  |  |  |
| --- | --- | --- | --- | --- | --- | --- | --- | --- | --- | --- | --- | --- | --- | --- |
| SDP | I | II | III | IV | V | X | Tot | I | II | III | IV | V | X | Tot |
| **12**<br>JU1511 & JU1926 vs.<br>JU1931 & JU1941 | 14 | 2 | 17 | 2 | 23 | 13 | 71 | 35 | 0 | 1 | 1 | 3 | 67 | 107 |
| **13**<br>JU1511 & JU1931 vs.<br>JU1926 & JU1941 | 6 | 1 | 14 | 39 | 41 | 3 | 104 | 0 | 2 | 106 | 13 | 15 | 27 | 163 |
| **14**<br>JU1511 & JU1941 vs.<br>JU1926 & JU1931 | 12 | 0 | 19 | 0 | 53 | 11 | 95 | 1373 | 0 | 119 | 5 | 103 | 44 | 1644 |
| **JU1511** | 37 | 32 | 61 | 14 | 18 | 81 | 243 | 457 | 28 | 211 | 20 | 9 | 430 | 1155 |
| **JU1926** | 0 | 32 | 4 | 59 | 5 | 1 | 101 | 5 | 44 | 5 | 26 | 10 | 5 | 95 |
| **JU1931** | 8 | 0 | 15 | 3 | 81 | 1 | 108 | 31 | 0 | 12 | 21 | 919 | 2 | 985 |
| **JU1941** | 76 | 0 | 66 | 5 | 38 | 22 | 207 | 150 | 1 | 155 | 35 | 94 | 1271 | 1706 |
| **Total** | 153 | 67 | 196 | 122 | 259 | 132 | **929** | 2051 | 75 | 609 | 121 | 1153 | 1846 | **5855** |

**Figure 1:**
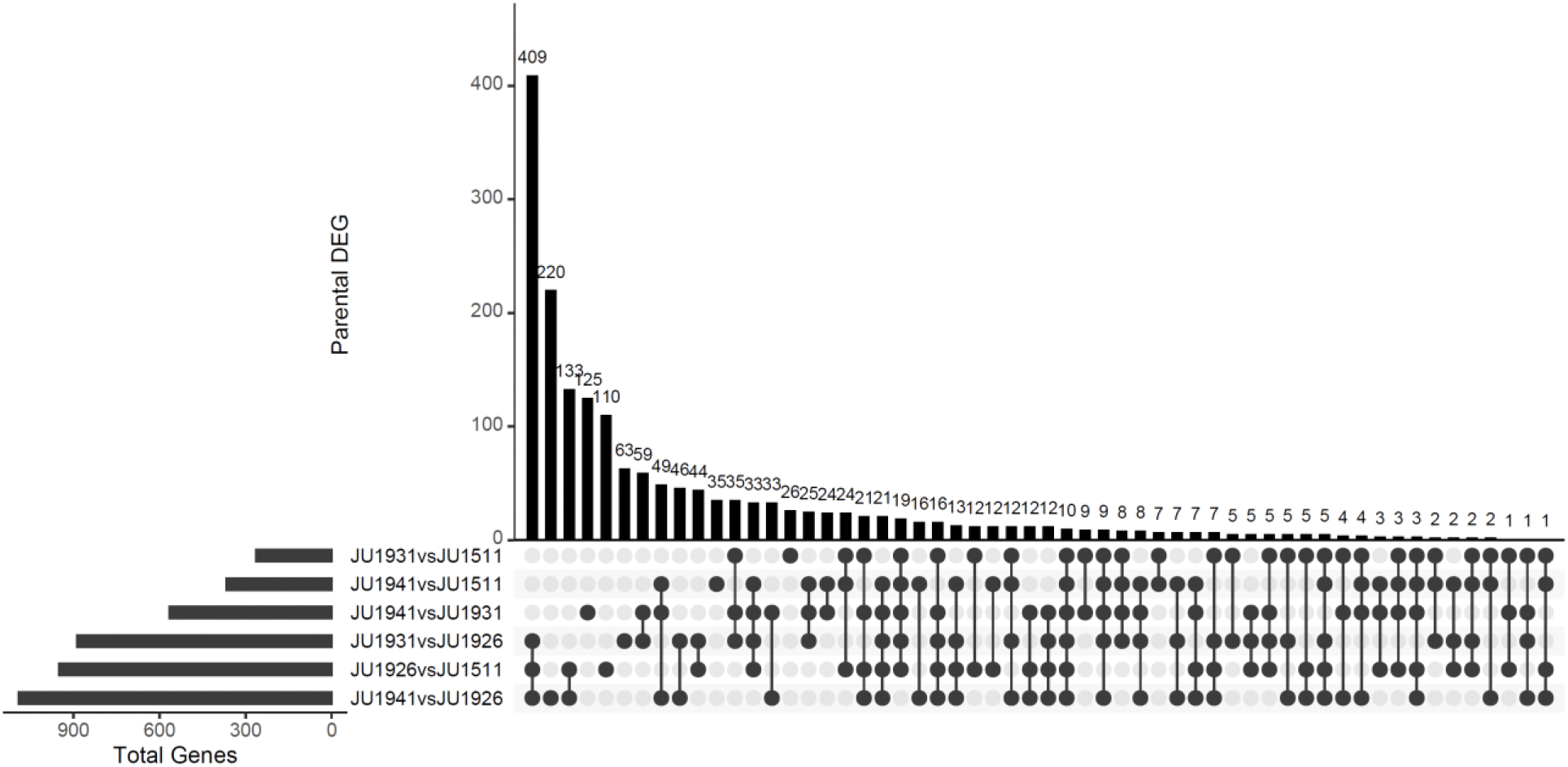
Gene expression differences between the four mpRIL parental lines. Upset plot shows the pairwise comparisons and the overlap between the pairs (TukeyHSD; p<0.001; FDR=0.05). The horizontal bar plot shows the number of differentially expressed genes per parental pair, while the vertical bar plot indicates the number of shared differentially expressed genes per comparison. For example, an overlap of 409 genes was found between the three comparisons that include the JU1926 parental line, which shows that JU1926 differed most from all other lines.

### Transgression and Heritability

To explore the variation in gene expression between the different parental and mpRIL genotypes we applied principal component analysis on the log_2_ gene expression ratios (**Figure2A**). Here we can see that the expression variation in many of the mpRILs extends beyond the parental expression variation, which suggests transgression. We quantified this and found transgression for 7,896 genes (FDR = 0.08; **Figure2B-C, Supplement table 2**). Notably, most transgression was one-sided, showing increased expression level beyond the highest expression level found in the parental lines. This suggests that multiple segregated loci are involved in regulating the transcription in the mpRILs. Transgression often indicates that the trait variation, in this case gene expression levels, is heritable. We calculated the narrow sense heritability (*h*^2^) and found significant *h*^*2*^ for expression variation of 9,500 genes (FDR = 0.05; **Figure 2D, Supplement table 2**). Most gene expression variation showed a *h*^*2*^ below 0.5, indicating that part of the variation is caused by other factors than additive genetic effects. These factors could be technical, environmental but also more complex genetic interactions such as epistasis.

**Figure 2:**
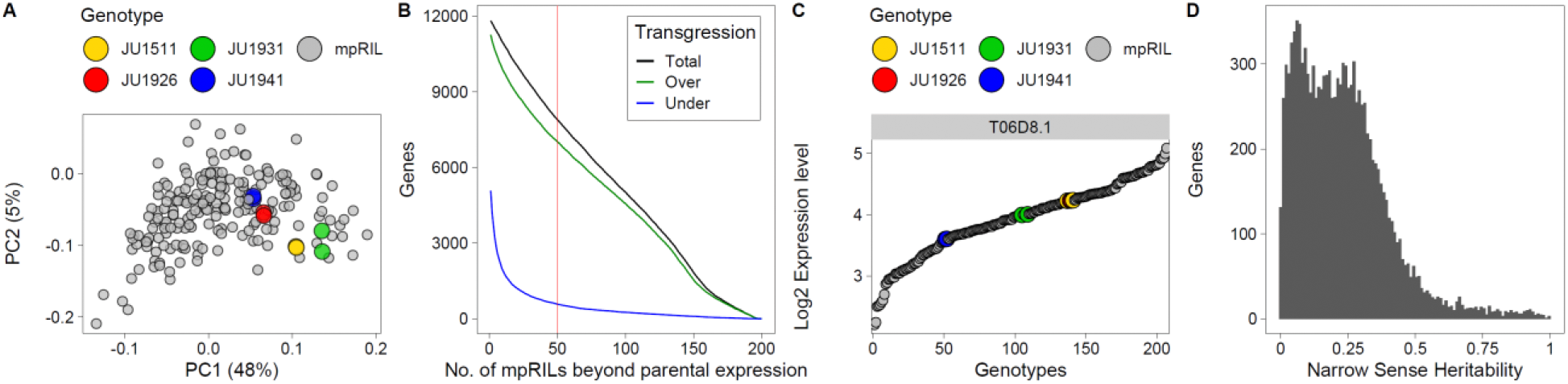
Gene expression variation in the mpRILs and parental genotypes. **A)** Principal component analysis (PCoA) of the log_2_ ratios, mpRILs shown in grey, parental lines shown in colour. **B)** Transgression: number of mpRILs beyond the parental expression level (x-axis) against the number of genes (y-axis). The mpRILs below (under) the lowest parental expression level in blue, mpRILs over the highest parental expression level in green and the sum of these (total) in black. **C)** Example of two-sided transgression for expression levels of gene T06D8.1. **D)** Genes with significant narrow sense heritability (*h*^*2*^) and the distribution of heritable variation of gene expression variation at FDR = 0.05.

### Expression QTLs

To find the loci involved in gene expression variation between the mpRILs we used a single marker QTL model. We found 6,784 eQTLs (one eQTL per gene, -log_10_(p) > 5.35; FDR = 0.01), of which 929 were *cis-*and 5,855 *trans*-eQTLs (**Table 1**; **Figure 3; Supplement table 2**). Most *cis*-eQTLs were found on chromosome V and most *trans*-eQTLs on chromosomes I and X. For both *cis*- and *trans*-eQTLs, fewest where found on chromosome II and IV. The SNP Distribution Pattern (SDP) groups SNPs with the same distribution in the parental lines. When the SDP is considered, many of the *cis*-eQTLs were found to have an effect where either the JU1511 or JU1941 allele was different from the three other parental genotypes. For the *trans*-eQTLs the largest groups also show this allelic difference or those SNPs that distinguish JU1511/JU1941 from JU1926/JU1931. A substantial group was found for the JU1931 allele, whereas hardly any were found for the JU1926 specific SNPs. The lack of JU1926 is somewhat surprising as it had the most differentially expressed genes (DEG) in the comparison of the parental lines, however we found much more genes with eQTLs than being DEG in the parental comparison. These are much more likely to be caused by new allelic combinations present in the mpRILs. Overall, the majority of the eQTLs are found on a few major effect loci with a specific SDP linkage (**Figure 3**). Moreover, comparing the *h*^*2*^ to the eQTLs showed that genes with an eQTL have a much higher *h*^*2*^ than those without an eQTL, where genes with an *h*^*2*^ > 0.25 almost all have an eQTL (**Figure 4**). Comparing *cis*- and *trans-*eQTLs showed that genes with a *cis*-eQTL have a higher *h*^*2*^ on average, yet the *h*^*2*^ distributions of *cis*- and *trans*-eQTLs are overlapping.

**Figure 3:**
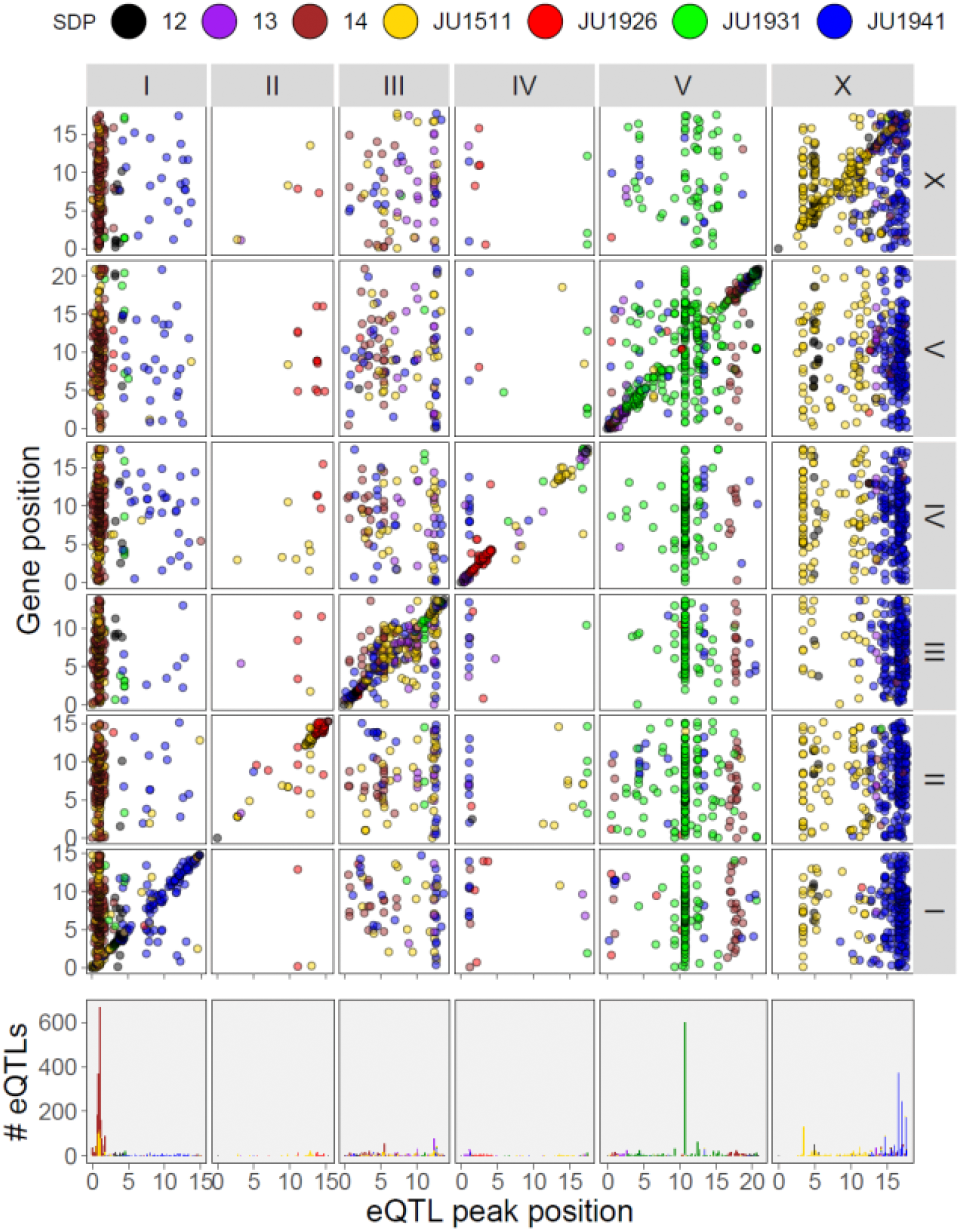
*Cis*/*Trans* plot of the identified eQTLs. eQTL position shown on the x-axis, gene position shown on the y-axis (upper plot) or number of eQTLs (bottom plot). SDP shown in colour, chromosomes shown in the grey strips on top and on the right of the panels.

**Figure 4:**
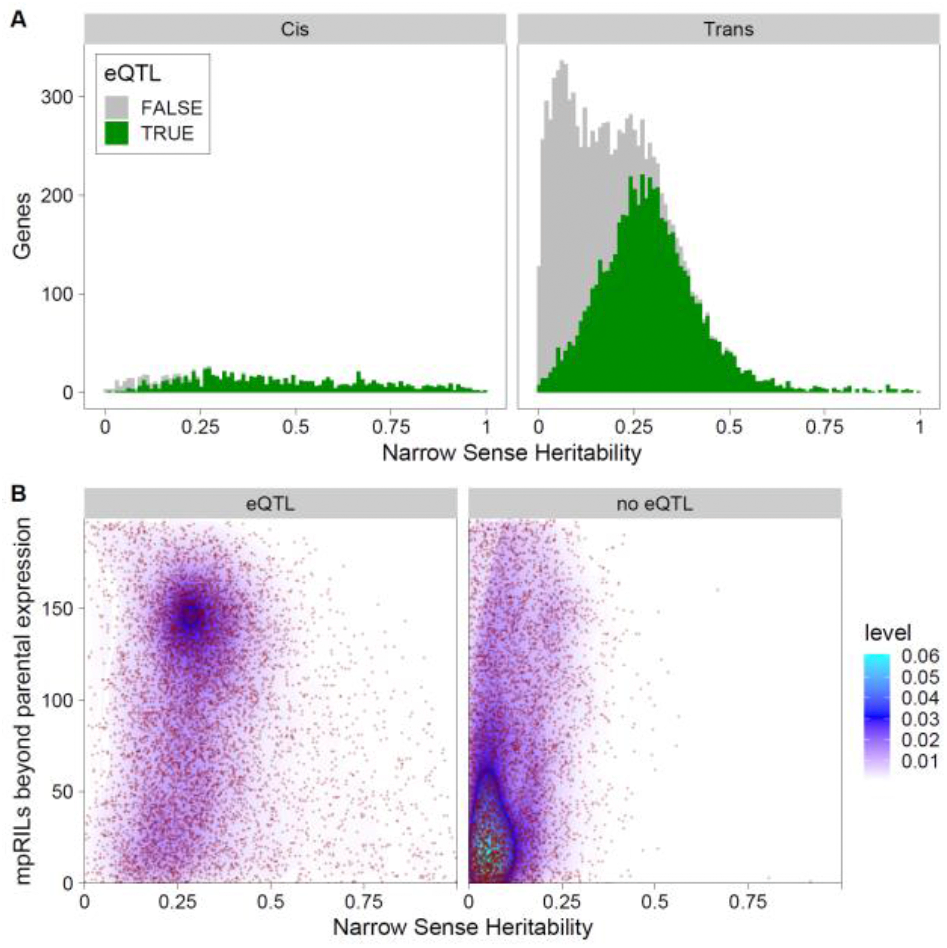
Relation between eQTLs, transgression and Narrow Sense Heritability (*h*^*2*^). **A)** Narrow Sense Heritability (*h*^*2*^; x-axis), distribution in genes (y-axis) with *cis-* and *trans*-eQTLs, significance of the eQTLs is TRUE (green) when > 5.35 and FALSE (grey) otherwise. **B)** Relation between Narrow Sense Heritability (*h*^*2*^; x-axis) and transgression (y-axis) for genes with and without a significant eQTL, individual datapoints shown in red, colour gradient indicates datapoint density.

### Trans-bands

A large majority of the *trans*-eQTLs (3,704; 63% of all *trans*-eQTLs) were found in six hotspots, so called *trans*-bands (TBs) (number of *trans*-eQTLs > 100, window 1Mbp to both sides; **Table 2**; **Figure 3**). Two TBs were found on chromosome I, one on chromosome V and three on chromosome X. The two TBs on chromosome I co-located but were linked to different SDP: the SDP 14 (JU1511/JU1941 vs JU1926/JU1931) and SDP JU1511 (*vs*. the rest). The TB on chromosome V was linked to SDP JU1931 and the three TBs found on chromosome X were linked to SPD JU1511 and JU1941.

**Table 2:** Descriptive overview of the 6 identified trans-bands. SNP Distribution Pattern (SDP), Chromosome, Peak position and left and right borders in Mega-base pairs. Selection of enriched GO terms from supplement table 3 and overlap with phenotypic QTLs found in Snoek *et al* 2019 [45].

|  | SDP | CHR | Peak (Mbp) | Left (Mbp) | Right (Mbp) | eQTLs | GO Enrichment (selection from enrichment table) | Phenotypic QTL (in Snoek *et al.* 2019 [45]) |
| --- | --- | --- | --- | --- | --- | --- | --- | --- |
| **TB1** | 14<br>(JU1511 & JU1941 vs JU1926 & JU1931) | I | 1.03 | 0.03 | 2.03 | 1339 | thermosensory behaviour, negative regulation of engulfment of apoptotic cell, DNA replication, embryonic body morphogenesis, establishment or maintenance of actin cytoskeleton polarity, muscle fiber development, epidermis development, response to unfolded protein and, molting cycle, collagen and cuticulin-based cuticle, | Population growth on *Erwinia* and on *B. thuringiensis* |
| **TB2** | JU1511 | I | 0.83 | 0 | 1.83 | 443 | regulation of protein stability, regulation of vulval development, DNA replication, anaphase-promoting complex and, microtubule polymerization | NA |
| **TB3** | JU1931 | V | 10.74 | 9.74 | 11.74 | 607 | hemidesmosome assembly, external side of plasma membrane and, negative regulation of response to oxidative stress | NA |
| **TB4** | JU1511 | X | 3.40 | 2.40 | 4.40 | 133 | few | heat-shock sensitivity |
| **TB5** | JU1941 | X | 14.69 | 13.69 | 15.64 | 225 | few | population growth on *B. thuringiensis* |
| **TB6** | JU1941 | X | 16.60 | 15.65 | 17.6 | 957 | embryonic body morphogenesis, DNA replication, integral component of peroxisomal membrane, anaphase-promoting complex, endosome, phagocytic vesicle membrane, neuronal signal transduction, response to anoxia, cuticle pattern formation, cell fate commitment, hemidesmosome associated protein complex and, response to lipid | sensitivity to oxidative stress |

### GO enrichment

To study the effect of TBs on biological function we used GO term enrichment (**Table 2, Supplement table 3**). Each of the TBs was linked to mostly different sets of GO terms, suggesting an effect on different parts of *C. elegans* biology. The genes mapping to TB1 on chromosome I were enriched for behaviour and muscle and epidermis development GO categories. The genes mapping to TB2 on chromosome I were enriched for the GO term “vulval development”, among others. The genes with a *trans*-eQTL on TB3 on chromosome V were enriched for GO terms associated with oxidative stress. The genes mapping to TB4 and TB5 on chromosome X only showed a few GO terms to be enriched and the genes mapping to TB6 on chromosome X were enriched for the GO term “response to anoxia” and many more. This shows that these TBs can be involved in several developmental processes and in the interaction with the environment.

### Overlap with other eQTL experiments

To investigate if the genes with eQTLs found in the present mpRIL study also had eQTLs in other studies, we compared them with the studies found in WormQTL2 (**Table 3**; [26, 28, 30, 31, 37, 39, 56]). In general, we found that a substantial group of genes with a trans eQTL in any of the studies had an eQTL in our mpRIL experiment (26.5% - 36.9%). The groups of genes with *trans*-eQTLs show much higher overlap than the genes with a *cis*-eQTL in any of the experiments (10.2% - 20.0%). Around a third of the genes with a *trans*-eQTL in Vinuela & Snoek *et al*. 2010 and Snoek & Sterken *et al*. 2017 also showed a *trans*-eQTL in the mpRILs, with numbers almost equal between developmental stages and treatments. Slightly fewer overlapping genes with eQTLs were found with Rockman *et al*. 2010 and Sterken *et al*. 2017. Comparing the experiments performed with the same N2 x CB4856 in the same lab, Li *et al* 2006, Vinuela & Snoek *et al*. 2010, Snoek & Sterken *et al* 2017, shows that environmental conditions and developmental stage only have a small effect on the global overlap and difference between *cis*-and *trans*-eQTLs. As the genetic backgrounds of the mpRILs are different from the N2 x CB4856 populations used in the other experiments, the low percentage of overlapping *cis*-eQTLs could be expected. The large group of genes with a *trans*- eQTL in both experiments shows that the expression levels of a substantial group of genes are more prone to be affected by genetic variation independent of environment or developmental stage, while the loci involved are most likely different in each experiment/condition [28, 30, 31].

## Discussion

In this experiment we used a population of multi-parental RILs (mpRILs) and RNA-seq to find 6,784 expression quantitative trait loci (eQTLs), of which 929 were *cis*-eQTLs and 5,855 were *trans*-eQTLs. A large proportion (63%) of the *trans*-eQTLs were found in six *trans*-bands. The total number of eQTLs found in this mpRIL study (6,784) is at the high end of what was previously found in other experiments (mean: 2,560; 653 – 6,518) [28, 30, 31, 37, 39]. This number is hard to compare as the number of identified eQTLs depend on many factors, such as population size, number of recombinations, statistical model, and RNA measurement technology used, which are nearly all different between this and the other eQTL studies in *C. elegans* [28, 30, 31, 37, 39]. Nevertheless, it seems that a combination of RNA-seq and multiple genetic backgrounds increased the number of detected eQTLs. A very clear increase was found for *trans*-eQTLs (5,855) compared to the numbers found in previous studies, even at a much lower significance threshold. For example, the study of Rockman *et al*. 2010 used a comparable number of recombinant inbred advanced intercross lines (RIAILs) as the number of mpRILs in this study (∼200), yet found fewer *trans*-eQTLs, however the different conditions and technologies used prevent any definitive conclusions. With respect to *trans*-eQTLs we do know that they depend on environmental conditions or a response to changing conditions. It could be that with a background of four parental genotypes the mpRILs perceive the ambient environment in a broader range than the RIAILs with a background of two parental genotypes used by Rockman *et al*. 2010, and the RILs in the other studies. For example, the mpRILs could have inherited parts of four different sets of environmental preferences as opposed to two in the RIAILs and RILs, potentially extending the accompanying gene expression patterns and eQTLs. Yet, the most likely reason for the increased number of *trans*-eQTLs is the use of RNA-seq in this study compared to micro-arrays in the other studies. Another reason for finding more *trans*-eQTLs could be due to the generally genome-wide equal allelic distributions in this population [45]. Namely, a similar *trans-*band as the chromosome I *trans*-band at 1 Mb (TB1) related to development has been spotted before, but has been spurious due to the *peel-1 zeel-1* incompatibility near that location [16, 28, 39]. Another advantage of using RNA-seq is that the genotype and gene-expression levels can be obtained from the same sample, preventing mis-labelling errors and the need for “reGenotyping” [64]. In summary, as has been shown for yeast [65], the combination of generally smaller effect of *trans*-eQTLs and higher dynamic range of RNA-seq would at least increase the possibility to pick-up *trans*-eQTLs in *C. elegans* and in general.

**Table 3:** **Overlapping eQTLs** between this mpRIL experiment and the RIL experiments available in WormQTL2 [63]. Percentages indicate the percentage of eQTLs found in the indicated experiment that are also found in the mpRILs eQTLs. Threshold used for the eQTL experiments in this table: -log10(p) > 3.5; *Cis*-eQTLs were called if the peak of the eQTL was within 1Mbp of the gene start, otherwise it was called a *trans*-eQTL.

| eQTL experiment | Total<br>*Cis* | *Cis*<br>Overlap(%) | Total<br>*Trans* | *Trans*<br>Overlap(%) |
| --- | --- | --- | --- | --- |
| Li *et al.* 2006 16°C [37] | 240 | 14.6 | 817 | 31.6 |
| Li *et al.* 2006 24°C [37] | 337 | 12.2 | 998 | 30.5 |
| Li *et al.* 2010 [26] | 752 | 14.5 | 3544 | 28.7 |
| Rockman *et al.* 2010 [39] | 1958 | 12.0 | 2792 | 28.8 |
| Snoek & Sterken *et al.* 2017 control [28] | 961 | 17.1 | 1481 | 36.1 |
| Snoek & Sterken *et al.* 2017 heat-shock [28] | 976 | 20.0 | 2776 | 36.9 |
| Snoek & Sterken *et al.* 2017 recovery [28] | 992 | 16.1 | 1519 | 33.4 |
| Sterken *et al.* 2017 [30] | 719 | 10.2 | 1116 | 26.5 |
| Vinuela & Snoek *et al.* 2010 juvenile [31] | 303 | 11.9 | 2206 | 33.4 |
| Vinuela & Snoek *et al.* 2010 old [31] | 220 | 15.0 | 1790 | 34.9 |
| Vinuela & Snoek *et al.* 2010 reproductive [31] | 348 | 13.2 | 2010 | 32.7 |

We previously found QTLs for several different phenotypes, such as population growth on different bacteria, sensitivity to heat-shock and oxidative stress [45]. Four *trans*-bands were found to co-locate with the previously found phenotypic QTLs (**Table 2**). Population growth on *Erwinia* and on *B. thuringiensis* DSM was found to co-locate with TB1, which was enriched for GO terms related to muscle, epidermis, and moulting. This could indicate a difference in these structures that can affect the interaction with different types of bacteria or could indicate that there is a difference in developmental speed through which differences in the expression, and subsequent eQTLs, of moulting related genes are picked up. A QTL for heat-shock sensitivity was inferred to co-locate with TB4, however no indication for a link with this phenotype was found in the annotation of the genes with an eQTL at this position. The same was observed for TB5 and the overlap with population growth on *B. thuringiensis*, where GO enrichment also did not provide any leads to a potential mechanistic link. The overlap between the QTL for sensitivity to oxidative stress and TB6 however did show some clues from GO enrichment as genes involved in the peroxisome as well as DNA replication and cuticle formation could be involved in dealing with oxidative stress.

We expect to have only found a fraction of the eQTLs, as we only used a simple additive mapping model, a conservative score of one eQTL per gene, and standard lab conditions with only one time point for RNA isolation. Both the number of eQTLs and genes with one or more eQTLs are expected to increase when more complex models are applied to this data and/or different experimental conditions and time points are considered. Moreover, we use a SNP-based method for eQTL mapping, which has a binary option for each marker and therefore does not consider the genetic origin (parent) of the SNP. Using the genetic origin of the SNPs could reveal the more complex genetic interactions that could underly the differences in transcript levels between the mpRILs. These complex genetic interactions are suggested to be present in this mpRIL population, by the heritability and transgression found. A model in which each marker has the four parental options might indicate loci with more than two alleles affecting gene expression. Furthermore, some (relatively small) genetic loci might have been missed all together as our investigations are based on the N2 reference genome and wild-isolates can have vastly divergent regions of which sequences reads fail to align to the N2 reference genome with conventional methods [49].

This study provides a more detailed insight into the genetic architecture of heritable gene expression variation in a multiparent recombinant inbred population. The use of RNA-seq data in combination with more than two alleles allows for a more precise detection of QTLs and incorporates a wider band of standing genetic variation, resulting in a substantial increase in eQTLs especially *trans*-eQTLs. Comparison to bi-allelic studies supports the position of eQTLs and may be used to detect a more detailed pattern of associated loci. We expect this study, data, and results to provide new insights into *C. elegans* genetics and eQTLs in general as well as to be a starting point to further test and develop advanced statistical models for detection of eQTLs and systems genetics studying the genotype-phenotype relationship.

## Methods

### Nematode strains and culturing, RNA-sequencing, Construction of the genetic map

The *C. elegans* strains and culturing condition, RNA-sequencing and construction of the genetic map can all be found in Snoek *et al*. 2019. RILs, Genetic map and eQTL profiles can found on WormQTL2 [66] (http://www.bioinformatics.nl/EleQTL; Snoek & Sterken *et al*. 2020 [56])

### SNP calling and gene expression levels

The paired end reads were mapped against the N2 reference genome (WS220) using Tophat [67], allowing for 4 read mismatches, and a read edit distance of 4. SNPs were called using samtools [66], mpileup with bcftools and vcfutils also described in Snoek *et al*. 2019 [45]. Expression levels were determined using the tuxedo pipeline [68]. Transcripts were assembled from the mapped reads using cufflinks [68]. Raw RNA-seq data can be found in the Sequence Read Archive (SRA; https://www.ncbi.nlm.nih.gov/sra) with ID PRJNA495983. Normalized read-counts can be found on WormQTL2 (http://www.bioinformatics.nl/EleQTL; [56])

### Heritability and Transgression

Heritability of gene expression levels was calculated using the heritability package in “R”. A narrow-sense heritability was calculated using the function *marker_h2* [69]. The required kinship matrix was calculated using the *emma*.*kinship* function from the EMMA package [70]. To determine significance, we used a permutation approach where we shuffled the expression levels per transcript. After 100 permutations, the 95^th^ highest value was taken as the 0.05 false-discovery rate [69, 71, 72]. Transgression was determined by counting the number of mpRILs with an expression level beyond the mean + 2SD of the most extreme parental lines. SD was calculated on the within variation of the parental samples. False discovery rate (FDR) was determined by permutations, randomly assigning the parental labels to gene-expression values.

Transgression was evaluated at an arbitrary 50 mpRILs (25% of all lines; FDR = 0.08) beyond the most extreme parental lines.

### eQTL mapping and FDR

For eQTL mapping we first selected the genes with consistently found transcripts, meaning those expressed in at least 20 samples with a mean log_2_ expression (fpkm) > -5. eQTLs were mapped by a linear model using a single maker model explaining gene expression (as log_2_ ratio with the mean) by one SNP-marker at the time for the whole genome. False Discovery Rate (FDR) was determined by one round of permutations where for each transcript the counts were randomly distributed over the RILs before eQTL mapping. The - log_10_(p) value when number of false positives divided by the number of true positives was < 0.01 (-log_10_(p) > 5.35). Genome wide eQTL significance profiles (-log10(p)) can be found on WormQTL2 (http://www.bioinformatics.nl/EleQTL; [56])

### Enrichment analysis and figures

Enrichment of GO terms was done using the hyper-geometric test in “R” [73]. GO term genes associations were download from Wormbase (www.wormbase.org) version WS276. Only genes that passed the filtering step for eQTL mapping where used as background genes. For significant enrichment, a p-value < 1e^-5^ was used and a geneset size per GO term > 3. Most figures were made using the R package ggplot2 [74] except figure 1 which was made using the UpSetR library.

### eQTL comparison between experiments/studies

To compare how many genes with an eQTL overlapped between the different studies [26, 28, 30, 31, 37, 39, 56] available in WormQTL2 [56], we downloaded the eQTL profiles and markers used per experiment and listed the genes with a *cis* or a *trans* eQTL. For eQTL determination, the most significant marker per gene was taken as the peak. A -log_10_(p) > 3.5 was use as threshold for calling the eQTL. An eQTL was determined *cis* when the peak position was within 1Mbp of the start position of the gene. These lists were compared with the genes having an eQTL in this study. The percentage overlap was calculated against the original study.

## Supporting information

Supplement_tables_Snoek_etal_2021

## Acknowledgements

We acknowledge financial support from the Deutsche Forschungsgemeinschaft to HS (grant number SCHU 1415/11 and project A1 within the CRC 1182), and to PCR (Competence Centre for Genomic Analysis (CCGA) No. 07495230). JK was funded by NIH grant 1R01AA 026658. Furthermore, financial support from the NWO-ALW (project 855.01.151) to RJMV was given. MGS was supported by NWO domain Applied and Engineering Sciences VENI grant (17282). The funders had no role in study design, data collection and analysis, decision to publish, or preparation of the manuscript.

## Author contributions

LBS, HS and JEK conceived the study, RJMV and JR performed the experiments, PCR coordinated and supervised transcriptome sequencing, LBS, MGS and HN analysed the data, LBS wrote the paper with contributions from all authors.

